# Translational adaptation of human viruses to the tissues they infect

**DOI:** 10.1101/2020.04.06.027557

**Authors:** Xavier Hernandez-Alias, Martin H. Schaefer, Luis Serrano

**Affiliations:** Centre for Genomic Regulation (CRG), The Barcelona Institute of Science and Technology, Dr. Aiguader 88, Barcelona, 08003, Spain; IEO European Institute of Oncology IRCCS, Department of Experimental Oncology, Via Adamello 16, Milan, 20139, Italy; Universitat Pompeu Fabra (UPF), Barcelona, 08002, Spain; ICREA, Pg. Lluís Companys 23, Barcelona, 08010, Spain

**Keywords:** tropism, tissue, translation, tRNA, codon usage, SARS-CoV-2, coronavirus

## Abstract

Viruses need to hijack the translational machinery of the host cell for a productive infection to happen. However, given the dynamic landscape of tRNA pools among tissues, it is unclear whether different viruses infecting different tissues have adapted their codon usage toward their tropism. Here, we collect the coding sequences of over 500 human-infecting viruses and determine that tropism explains changes in codon usage. Using an *in silico* model of translational efficiency, we validate the correspondence of the viral codon usage with the translational machinery of their tropism. In particular, we propose that the improved translational adaptation to the upper respiratory airways of the pandemic agent SARS-CoV-2 coronavirus could enhance its transmissibility. Furthermore, this correspondence is specifically defined in early viral proteins, as upon infection cells undergo reprogramming of tRNA pools that favors the translation of late counterparts.

## INTRODUCTION

Given the degeneracy of the genetic code, multiple 3-letter combinations of nucleotides can code for the same amino acid. Such synonymous codons are nevertheless not uniformly distributed along the genomes and can significantly deviate between organisms^1^. Evolutionary forces that explain the existence of the so-called codon bias include (1) a mutation pressure for a certain GC base composition depending on the species and chromosomal location, and (2) the translational selection for codons corresponding to highly expressed tRNA isoacceptors^2–4^.

Viruses strongly depend on the translational machinery of the host for the expression of their own proteins and, ultimately, their replication. For instance, given the small size of most viral genomes, no or very few tRNA genes are generally autonomously encoded^5^. In terms of codon usage, it has indeed been shown that bacteriophages are specifically adapted to their microbial hosts^6,7^. This information has been applied in the prediction of viral hosts from metagenomics data^8,9^. The codon usage of human-infecting viruses is similarly adapted to the host^10,11^, and actually the concept of codon deoptimization has been applied in the design of attenuated vaccines^12^.

Although translational selection has long been under debate in human^13^, recent studies indicate that different tissues and conditions showcase distinct tRNA expression profiles, leading to changes in their respective translational efficiency^2,14^. In agreement with this observation, the codon usage of papillomavirus capsid proteins is adapted to the tRNAs of differentiated keratinocytes, where their translation becomes specifically efficient^15,16^. In addition, upon HIV-1 infection, the host tRNA pool is reprogrammed to favor translation of late viral genes^17^, a phenomenon that is indeed exploited by host antiviral mechanisms^18^. Furthermore, some viruses with a specific tissue tropism resemble the codon bias of highly expressed proteins of their respective infecting tissues^19^. Nevertheless, despite the few aforementioned studies, a high-throughput analysis of the translational selection of viral genomes to their tissue tropism has been heretofore hindered by the absence of tissue-wide tRNA expression data.

Here, we systematically analyze the relative codon usage landscape of over 500 human-infecting viruses together with the recently published tRNA expression profiles of human tissues^2^. Among other viral annotated features, including phylogeny and Baltimore classification, their tissue tropism explains more variance in codon usage than the other tested features. In consequence, tropism corresponds with codon optimization patterns that can be associated with tissue-specific profiles of tRNA-based translation efficiencies.

The SARS-CoV-2 coronavirus constitutes the etiologic agent of the biggest pandemic of the 21^st^ century, causing the COVID-19 pneumonia-like disease. Originated in Wuhan (China), the virus has already caused 40,598 deaths as of 1^st^ April 2020^20^. In this context, recent scientific efforts suggest that the novel coronavirus strain has evolved to preferentially infect nasal goblet and ciliated cells as well as the type-2 alveolar epithelial cells^21,22^. However, some non-respiratory-related symptoms such as loss of smell and taste hint as well to other tropism potential outside the respiratory tract^23,24^. In terms of translational efficiency, we here discover that the coronavirus proteome is especially adapted not only to upper respiratory airways and alveoli, but also to the gastrointestinal tract and brain.

Further, by studying the tissue-adaptation among the viral proteome, we also determine that early replication-related proteins are more translationally-adapted than the late structural counterparts. Overall, we observe a tropism-specific adaptation of the viral proteome to the tRNA profiles of their target tissues, which opens the door to the development of tissue-specific codon-deoptimized vaccines and targeted antiviral therapies.

## RESULTS

### Tropism corresponds with differences in Relative Codon Usage of human-infecting viruses

Publicly available genomic data comprised a total of 564 human-infecting viruses, distributed among 33 families and covering all seven Baltimore categories (Sup. Table 1). Across this diversity, six main viral tropisms were defined for 182 viruses based on the ViralZone curated database^25^: neurons, immune cells, respiratory tract, hepatocytes, intestine, and epithelial cells (Fig. 1A). Their corresponding coding sequences constituted a total of 4935 viral proteins (Sup. Table 1), for which we determined the Relative Codon Usage (RCU, i.e. the contribution of each synonymous codon to the amino acid it encodes, see Methods).

**Figure 1.**
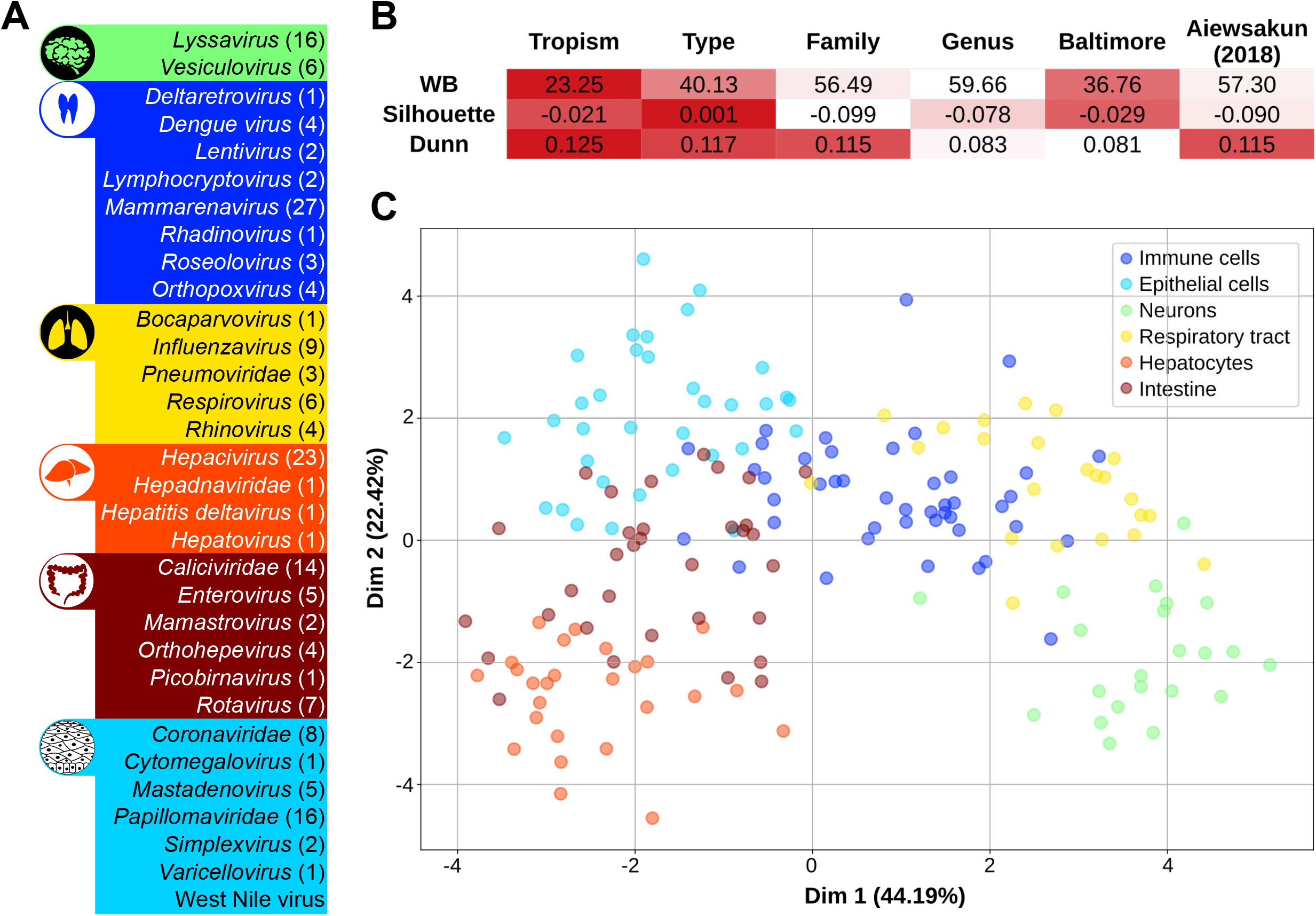
Tropism corresponds with differences in relative codon usage of human-infecting viruses. (A) A total of 564 viruses was distributed among 33 families and covered all seven Baltimore groups. From there, 182 viruses were classified in six general tropisms based on ViralZone annotations^25^. (B) Three internal clustering indexes were computed to assess the validity of each viral classification in terms of their RCU. Good cluster performances lead to low WB indexes, but to high Silhouette and Dunn values (as shown in the color code). (C) Linear Discriminant Analysis of the RCU of the 182 tropism-defined viruses. In brackets, the percentage of variance explained by each of the components.

In order to understand the main factors driving differences between viral RCU, we used three internal clustering indexes that assess how similar each virus is to a certain group compared to other groups. Taking the average RCU over each of the 564 viral proteomes, we applied this framework to assess the grouping performance of six different viral features: tropism, type of genetic material, family, genus, Baltimore category, and a sequence-based classification by Aiewsakun and Simmonds (2018). In such analysis, the tropism leads the best classification of viral RCUs, followed by the viral genetic type (Fig. 1B). On the other hand, classical and sequence-based phylogenetic classifications show poor clustering performances.

Given the impact of viral tropism on the RCU, we sought to determine the main codon differences between the six defined target tissues. By using a linear discriminant analysis (see Methods), the 182 tropism-defined viruses were classified in six clear clusters, regardless of other factors such as the phylogenetic lineage (Fig. 1C). In consequence, we observe that specific codon usage profiles are associated with the tissue tropism of human-infecting viruses.

### Viruses are adapted to the tRNA-based translational efficiencies of their target tissues

Based on the RCU differences between viruses with distinct tropism, we hypothesize that distinct tissues impose selection towards a certain set of translationally-efficient codons. However, a validation for this hypothesis requires the accurate quantification of tissue-specific tRNA profiles, which has been hitherto missing. With the advent of such high-throughput expression data^27,28^, here we retrieved the previously-published Supply-to-Demand Adaptation (SDA) estimate for translational efficiency^2^, which computes the balance between the supply (i.e., the anticodon tRNA abundances) and demand (i.e., the codons expressed in mRNAs) of each codon (see Methods).

Using a total of 620 healthy samples from The Cancer Genome Atlas (TCGA) dataset^2^, we first computed the SDA of all viral-protein-coding sequences based on the SDA weights of their constituent codons. Therefore, taking the average of all healthy samples across each of the 23 TCGA cancer types (Sup. Table 2), we determined the estimated translational efficiencies of viral proteins in different human tissues (Sup. Table 3).

Next, from the perspective of the translational selection hypothesis, we would expect that viral proteins are translationally adapted to their target tissues. In consequence, we tried to test our hypothesis using a completely blind and unbiased random forest classifier, which applies machine learning in order to predict the tropism of each viral protein based on the SDA to different tissues (see Methods). The resulting performance of the models, based on the Area Under the Curve (AUC) of their Receiver Operating Characteristic (ROC) curves, ranges between 0.74-0.91 (Fig. 2A), clearly higher than the no-skill model of 0.5. Similar results are also obtained from complementary prediction performance metrics such as Precision-Recall curves (Fig. 2A). These results indicate that our machine learning model is able to predict the tropism of a viral protein based on its SDA to tissues with high accuracy. In concordance, a linear discriminant analysis of the average SDA of each virus across tissues can similarly separate different clusters of viral tropism based on their translational efficiencies (Extended Data Fig. 1).

**Figure 2.**
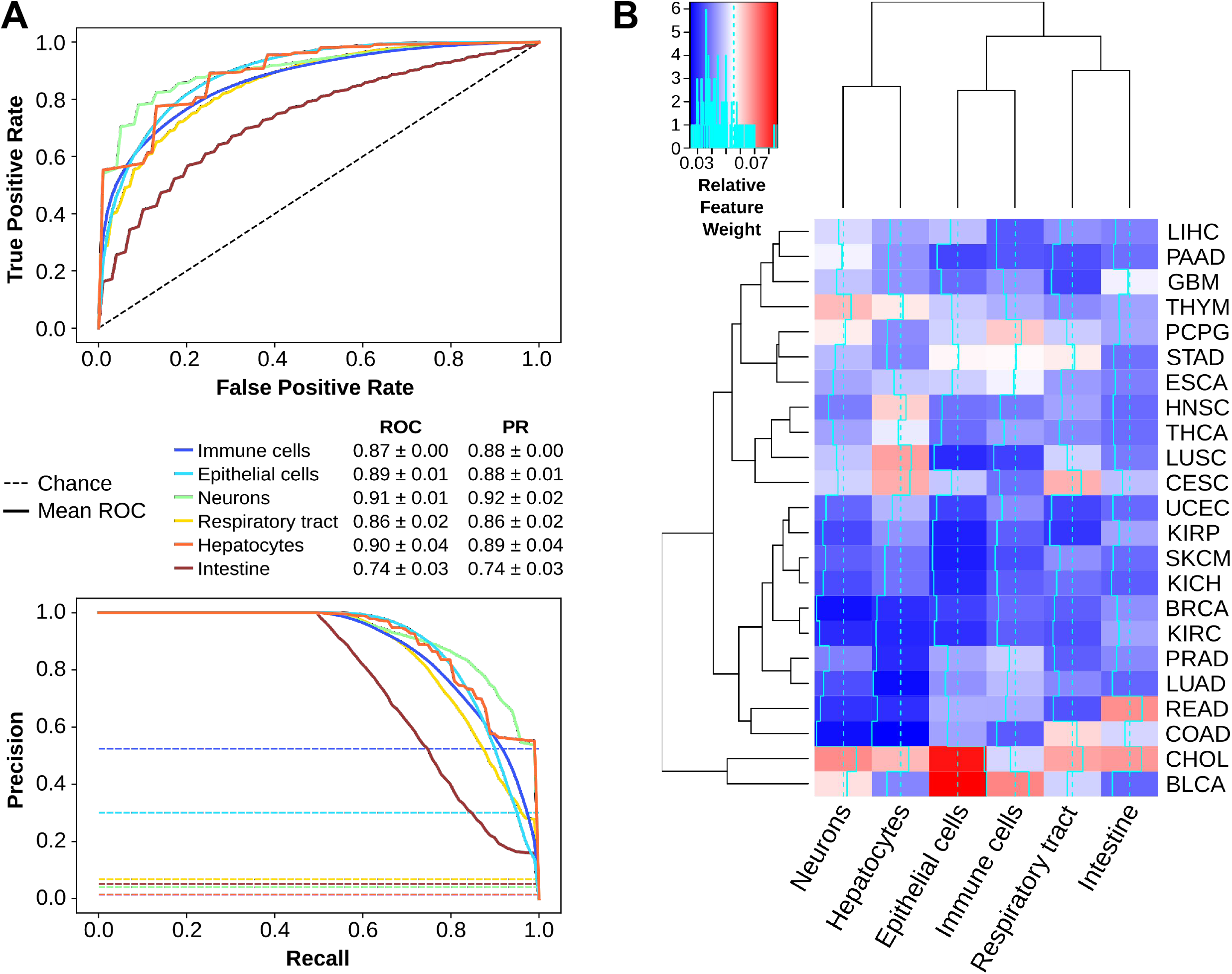
Viruses are adapted to the tRNA-based translational efficiencies of their target tissues. (A) Receiver Operating Characteristic (ROC) and Precision-Recall (PR) curves of a Random Forest Classifier, in which the average Supply-to-Demand Adaptation of viral proteins to each of the 23 TCGA tissues is used to predict their corresponding viral tropism (see Methods). The area under the curves (AUC) ± SD summarizes the performance of the model. (B) Relative feature weights of each of the 23 TCGA tissues for each of the six tropisms. The dendrograms show a hierarchical clustering among tissues (left) and among tropisms (top). Refer to Sup. Table 2 for full TCGA cancer type names. See also Extended Data Fig. 1.

In an attempt to understand which tissues are the most predictive in identifying the viral tropism of proteins, we analyzed the relative feature importance within each random forest classifier, which measures the contribution of each tissue SDA in the decision trees (Fig. 2B). The main observation is that no single tissue alone is able to discriminate against the specific tropism, since all feature importances lie below 0.10. However, it is also clear that translational adaptation to bile duct (CHOL, for healthy samples of cholangiocarcinoma) is a recurrent discriminant feature, while other tissues are specifically important for just one or few tropisms, such as rectum (READ, for healthy samples of rectum adenocarcinoma) in predicting intestinal viruses. In any case, the directionality of these features cannot be established.

Overall, as tropism of proteins can be predicted from their translational adaptation to tissues, these results indicate that viral proteomes are specifically adapted to certain tRNA-based translational efficiencies. In consequence, and complementary to the observations of mutational pressure driving viral codon bias^11,29,30^, we describe the basis for a potential tissuespecific translational selection of the viral codon usage.

### SARS-CoV-2 is translationally adapted to upper respiratory airways and alveoli

As our systematic analysis suggests that the codon usage of viruses tend to be adapted to the tissue they infect, we selected the novel coronavirus SARS-CoV-2 and other related respiratory viruses to further explore their translational adaptation profile over tissues.

We initially reconstructed tRNA expression profiles along the respiratory tract making use of the spatial information associated with healthy TCGA samples from head and neck squamous cell carcinoma (HNSC), lung squamous cell carcinoma (LUSC) and lung adenocarcinoma (LUAD) (Sup. Table 4). We then computed the SDA of viral proteins from the three pandemic coronaviruses of the last two decades SARS-CoV^31^, MERS-CoV^32^, and SARS-CoV-2^33^, as well as the common flu-causing influenza A virus (H1N1) along the respiratory tract. We find that the new coronavirus strain SARS-CoV-2 is the most translationally adapted virus across all tissues, especially in comparison with MERS-CoV and influenza A virus (Fig. 3A). Further, the novel coronavirus is highly adapted throughout the whole respiratory tract, and specifically the upper respiratory airways and the alveolar region in the lung periphery are the most efficient tissues.

**Figure 3.**
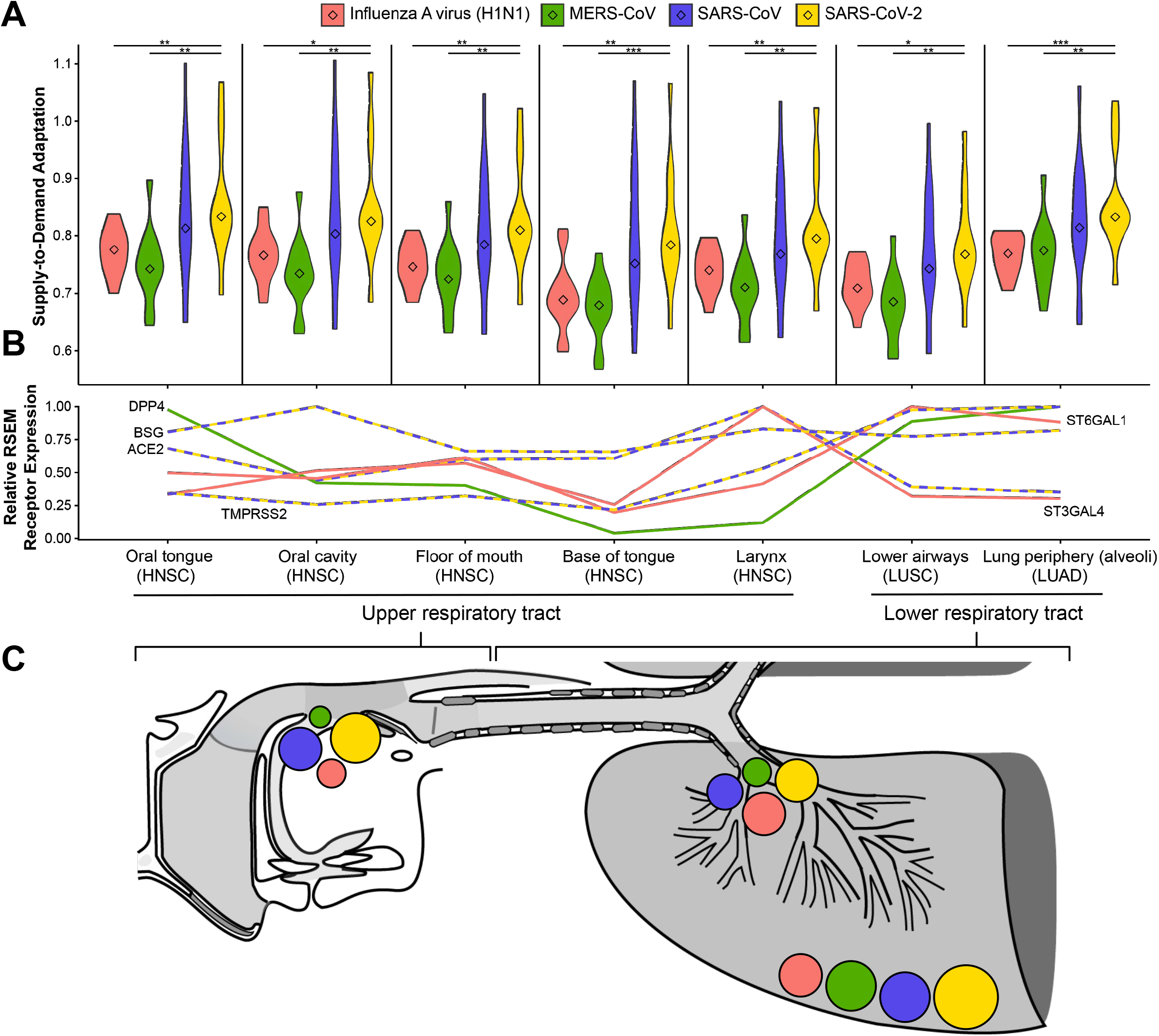
SARS-CoV-2 is translationally adapted to upper respiratory airways and alveoli. (A) Using the codon adaptation weights of all TCGA samples along the respiratory tract (HNSC, LUAD, LUSC), we compute the SDA of proteins from coronaviruses (and influenza A virus. Center values within the violin plot represent the median. Using a two-tailed Wilcoxon rank-sum test, only significant differences between viruses are shown: * (p <= 0.05), ** (p <= 0.01), *** (p <= 0.001), **** (p <= 0.0001). (B) Average RSEM expression of the corresponding entry receptors of the viruses: MERS-CoV binds to DPP4^35^, SARS-CoV and SARS-CoV-2 bind to ACE2 or BSG with priming by the protease TMPRSS2^36,37^, influenza A virus binds to sialic acids, which are synthesised by the enzymes ST3GAL4 and ST6GAL1^34^. RSEM receptor expression is re-scaled between 0 and 1 among all tissues, being 1 the tissue with highest expression. Refer to Sup. Table 3 for full TCGA cancer type names. (C) Schematic representation of the potential of infection of each virus based on the coincidence of both translational adaptation and entry receptor expression. See also Extended Data Fig. 2.

Moreover, given that the tropisms not only depend on the translational adaptation to the host, but also on the expression of the required entry receptors, we measure the respective receptors of each virus. Influenza A viruses bind to α(2,6)-linked and α(2,3)-linked sialic acids, which are synthesized by the enzymes ST6GAL1 and ST3GAL4, respectively^34^. While their expression is relatively uniform along the airways, the MERS-CoV uses the parenchyma-specific receptor DPP4 (Fig. 3B)^35^. In both cases, the expression of their entry receptor coincides with an optimal translational adaptation in the lower respiratory tract. On the other hand, both the SARS-CoV and SARS-CoV-2 strains bind to the ACE2 protein and require the proteolytic priming of the viral spike protein by TMPRSS2^36^, although the receptor BSG/CD147 has also been proposed^37^. In all cases, their expression throughout the airways (Fig. 3B) alongside their efficient translational adaptation defines the upper respiratory tract and the alveoli as their particular optimal tropism (Fig. 3C). This is in agreement with recent single-cell transcriptomic studies reporting the expression of ACE2 in the nasal goblet and ciliated cells as well as the type-2 alveolar epithelial cells^21,22^.

Apart from the clear viral tropism of SARS-CoV-2 to the respiratory tract, recent studies propose that their tropism can expand to other tissues such as the digestive system or the brain^23,24^. For this reason, we also extended our translational analysis to all the 23 tissues of the TCGA dataset (Extended Data Fig. 2A), together with the expression of the corresponding receptors (Sup. Table 4). In agreement with the clinical findings^23,24^, the gastrointestinal tract emerges as the most translationally adapted tissue, followed by the other epithelial-like tissues and the brain. Therefore, in terms of translational efficiency, the novel SARS-CoV-2 is widely adapted across tissues.

In an attempt to elucidate the translational selection that could have benefitted the evolution of the new coronavirus, we also compared the SDA adaptation of SARS-CoV-2 to those of close phylogenetic strains: the human-infecting SARS-CoV and the bat coronavirus RatG13, with 79.6% and 96.2% of sequence identity, respectively^33^. We observe that all four main proteins of the new strain have optimized their codon usage with regard to the previous 2003-pandemic SARS-CoV (Extended Data Fig. 2B). On the other hand, even though the ratio of synonymous versus nonsynonymous mutation rate of the bat coronavirus compared to SARS-CoV-2 is exceptionally high^38^, their proteins are very similarly adapted to most human tissues.

In short, we report that the new pandemic SARS-CoV-2 coronavirus has evolved to significantly increase its translational adaptation to the human tissues. While it might not be sufficient on its own, the coincidence of both translational efficiency and entry receptor expression supports an optimal tropism of the alveoli but also the upper respiratory tract (Fig. 3C), which has recently been related to the higher transmissibility of the novel strain^21^.

### Early viral proteins are better adapted and translationally deoptimized throughout the process of infection

Given the tropism-specific adaptation of viral RCU towards the translational machinery of tissues, we wondered whether certain genomic subsets were specifically adapted to the tissue of infection. In particular, we speculated that early replication-related proteins would further benefit from such adaptation than late structural proteins, since once the virus takes control of the cell it could change its tRNA expression program^17^.

To estimate the adaptation of each protein to the tRNA-based codon efficiencies of each tissue, we computed their SDA^2^ (Sup. Table 5). For that purpose, we matched each virus to the tRNAs of their tissues of infection (Sup. Table 5). In concordance with our hypothesis, based on current viral annotations (VOGdb, vogdb.org), we observed a small but highly significant shift in SDA between structural and replication proteins in all but hepatocyte-infecting viruses (Fig. 4A). However, the proteomes of this latter group are poorly annotated at the viral orthology database VOGdb, and separation between classes is often hindered by the polyprotein structure of their genomes^39^. Similarly, we performed a Gene Set Enrichment Analysis to identify which Virus Orthologous Groups (VOGs) were enriched in high-SDA or low-SDA proteins (Fig. 4B). As determined by current annotations^39^, top-VOGs mostly contained replication-related early proteins, whereas bottom-VOGs had structural late functions.

**Figure 4.**
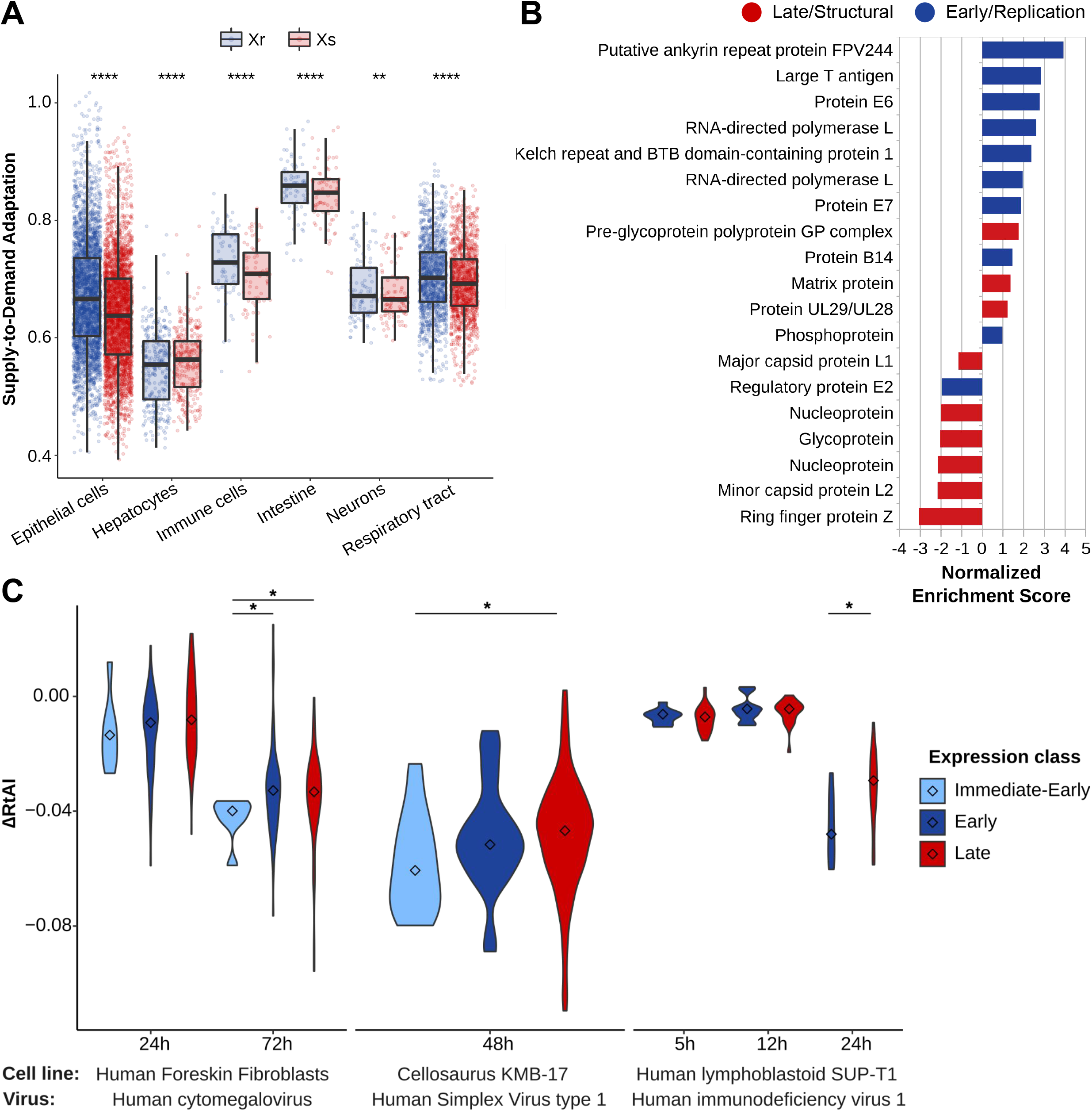
Early viral proteins are better adapted and translationally deoptimized throughout the process of infection. (A) Average Supply-to-Demand Adaptation of replication (Xr) and structural (Xr) proteins of a total of 104 annotated tropism-specific viruses, respectively matched to 461 samples of their tissues of infection (Sup. Table 5). Boxes expand from the first to the third quartile, with the center values indicating the median. The whiskers define a confidence interval of median ± 1.58*IQR/sqrt(n). Statistical significance is determined by paired (structural against replication proteins of each virus) and two-tailed Wilcoxon rank-sum test. (B) Top positive and negative Virus Orthologous Groups upon Gene Set Enrichment Analysis of the SDA of all proteins of tropism-specific viruses (Sup. Table 5). Based on their annotations, proteins groups are colored based on their early/replication or late/structural function^39^. (C) Differences of the relative tRNA Adaptation Index (see Methods) between mock and effective viral infections in different cell lines. Multiple time points are specified when available. Proteins are allocated to different time expression classes based on current viral knowledge^39^ (Sup. Table 6). Center values within the violin plot represent the median. Using a two-tailed Wilcoxon rank-sum test, only significant differences are shown: * (p <= 0.05), ** (p <= 0.01), *** (p <= 0.001), **** (p <= 0.0001).

Previous studies on the translational adaptation of the human immunodeficiency virus 1 suggested that the host tRNA pool is reprogramed upon viral infection in order to favor the expression of late genes^17^. In this direction, we wanted to test whether this tRNA reprogramming is a general adaptive mechanism among viral species. Using three previously published small RNA-sequencing datasets of human cell lines upon viral infection^40–42^, we quantified the tRNA of mock and productive infections at different time points (Sup. Table 6). Therefore, for all datasets, we detected a general decrease in translational efficiency upon viral infection, with late viral proteins being more favored than early ones specifically at longer infection times (Fig. 4C).

Overall, we determine that the tropism-specific adaptation of viruses is specifically pronounced among early proteins. However, later in infection, the tRNA pool of the host cells is reprogrammed to favor the expression of late viral genes, reproducing previous findings of HIV1 infection and extending the mechanism to dsDNA-based viruses such as human cytomegaloviruses and human simplex viruses.

## DISCUSSION

Tropism is determined by an ensemble of different factors, including the mechanism of viral entry to the host, the immune responses to the infection, or the viral hijacking of the cellular machinery in the interest of replication and propagation. In this article, we study the latter by focusing on the translational adaptation of viral genomes to the host.

While previous studies on the base composition and codon usage of both DNA and RNA viruses^11,30^ have attributed most of the codon usage variability to the mutational pressure of viral genomes, our analysis proposes tropism as an important driving force. By systematically interrogating all human-infecting viruses, we uncover that tissue tropism explains changes in their codon usage more than other viral properties such as type or family. Therefore, as mutational pressure would act more similarly within phylogenetically closer species, such tropism-related differences in codon usage suggests that tissue-specific tRNA expression could be driving a translational selection on viral genomes.

Although high-throughput sequencing of tRNAs has been only recently developed, cases of natural selection of codon usage towards the host have been previously proposed. For instance, codon usage of *Parvovirus* has been progressively adapted from dogs to cats after the host jump^43^. *Influenzaviruses* show a similar adaptation over time of viral isolation, deviating from the codon usage of avian hosts^44,45^. However, whether these progressive changes in codon usage over time are directly driven by translational selection has remained elusive. With the advent of tissue-wide datasets of tRNAs and their translational efficiencies^2^, we can now compute the Supply-to-Demand Adaptation (SDA) of all viral proteomes in different tissues. From there, we then created a random forest model that predicts with high accuracy the viral tropism of proteins based on their profile of adaptation to human tissues. In consequence, the tRNA-based adaptation profile of a protein is descriptive of their viral tropism, indicating that translational selection could indeed drive tropism differences of codon usage. It is important to remark that viruses could still have a good SDA to non-target tissues with similar tRNA expression patterns that are not infected because they are not exposed to the virus.

Specifically, we therefore focus our analysis on the new SARS-CoV-2 coronavirus, causative of the current COVID-19 pandemic. As has been recently suggested based on the expression of their entry receptor^21,22^, SARS-CoV-2 is also translationally adapted to the upper respiratory airways, as well as the lung parenchyma. This upper tract tropism has indeed been linked to the higher transmissibility of the strain^21^. The novel coronavirus showcases the highest SDA among all studied coronaviruses, including lung but also other tissues such as the digestive system and brain. Interestingly, recent reports have shown that COVID-19 patients very frequently have problems in the digestive tract^24,46,47^ as well as in neural tissue^23,48,49^.

One major open question in the field still persists: how much natural selection before or after zoonosis shapes the current coronavirus^50^. On the one hand, viruses tend to gradually change their codon usage after a zoonotic jump^43,45^. On the other hand, given the similarity of SARS-CoV-2 SDA with the phylogenetically closest bat coronavirus, it seems that a translational selection to increase SDA would have acted before the putative zoonosis from bats or other intermediate hosts. Furthermore, in agreement with the highest translational potential of SARS-CoV-2 in their target tissues, a recent model of viral tropism suggested that a tradeoff exists between the efficiency of viral translation and the translational load on the host, indicating that an improved codon usage can make the difference between symptomatic and natural hosts^51^.

On the other hand, it is known that host tRNA pools undergo reprogramming upon viral infection of HIV-1, vaccinia virus, and influenza A virus^17,52^. In this context, differences in codon usage between early and late viral genes have been previously reported, but the directionality of such changes remained unclear^10,53^. Based on our concept of tissue-specific adaptation, we therefore validate that early replication-related proteins are better adapted to the tissue of infection. Upon infection, we then unveil that changes in tRNA abundance switch the adaptation towards the expression of late structural proteins, confirming a general trend that had previously only been observed in HIV-1^17^.

Overall, this is the first systematic analysis establishing a link between the codon usage of human viruses and the translational efficiency of their tissue of infection. This correspondence is particularly observed in early viral proteins, while late counterparts benefit from the tRNA reprogramming that underlies the process of infection. We therefore envision the development of *ad hoc* gene therapies specifically targeting the tissue of interest.

## METHODS

### DATA SOURCES

#### Viruses and annotations

We included in the analysis all human-infecting viruses from the NCBI Viral Genome Browser, downloaded as of August 29, 2019. Additionally, for its interest, we added *a posteriori* the new SARS-CoV-2 virus. Viral metadata including family, genus, genetic material type and Baltimore category were retrieved either from the ICTV 2018b Master Species List^54^ or the ICTV Virus Metadata Resource (talk.ictvonline.org/taxonomy/vmr/). The sequence-based phylogenetic information was obtained from Aiewsakun and Simmonds (2018). Tissue and cell type tropism was determined based on the curated database ViralZone^25^, and allocated to each of the six main classes based on the main annotation. Sup. Table 1 contains all human-infecting viruses and their associated metadata.

#### Coding sequences

The coding sequences of human-infecting viruses from RefSeq were downloaded from the Codon/Codon Pair Usage Tables (CoCoPUTs) project release as of August 29, 2019^55,56^ (Sup. Table 1). The SARS-CoV-2 and RaTG13 sequences were directly downloaded from GenBank (Sup. Table 4).

#### Virus Orthologous Groups

Virus Orthologous Groups and their functional annotations (virus structure and replication) were downloaded from VOGdb (vogdb.org, release number vog94). The protein sets of each VOG were formatted to a Gene Matrix Transposed (GMT) file for custom GSEA analyses.

#### TCGA translational efficiency

The Supply-to-Demand Adaptation (SDA) is the balance between the supply (i.e. the anticodon tRNA abundances) and demand (i.e. the weighted codon usage based on the mRNA levels) for each of the 60 codons (excluding methionine and Stop codons)^2^. The SDA weights of all TCGA samples were downloaded from Synapse (www.synapse.org/tRNA_TCGA, syn20640275).

#### Small RNA-sequencing datasets upon viral infection

Three small RNA-sequencing datasets were downloaded to analyze the tRNA content of human cell lines upon viral infection. The non-productive mock infections of each dataset were also analyzed for normalization. In Stark et al. (2012), samples of Human Foreskin Fibroblasts (HFF) infected by human cytomegalovirus (HCMV) strain Towne at a multiplicity of infection (MOI) of 3, analyzed at 24 and 72 hours post-infection (GSE33584). In Shi et al. (2018), samples of cellosaurus KMB-17 infected by Human Simplex Virus type 1 (HSV1) strain 17 at a MOI of 1, analyzed at 48 hpi (GSE102470). In Chang et al. (2013), samples of lymphoblastoid SUP-T1 cells infected by Human immunodeficiency virus 1 (HIV1) strain LAI at a MOI of 5, at 5, 12 and 24 hpi (GSE57763).

### COMPUTATIONAL ANALYSIS

#### tRNA quantification

In small RNA-seq FASTQ files, sequencing adapters were trimmed using BBDuk from the BBMap toolkit [v38.22] (https://sourceforge.net/projects/bbmap): k-mer=10 (allowing 8 at the end of the read), Hamming distance=1, length=10-75bp, Phred>25. Using the human reference genome GRCh38 (Genome Reference Consortium Human Reference 38, GCA_000001405.15), a total of 856 nuclear tRNAs and 21 mitochondrial tRNAs were annotated with tRNAscan-SE [v2.0]^57^.

Trimmed FASTQ files were then mapped using a specific pipeline for tRNAs^2,58^. Summarizing, an artificial genome is first generated by masking all annotated tRNA genes and adding pre-tRNAs (i.e. tRNA genes with 3’ and 5’ genomic flanking regions) as extra chromosomes. Upon mapping to this artificial genome with Segemehl [v0.3.1]^59^, reads that map to the tRNA-masked chromosomes or to the tRNA flanking regions are filtered out in order to remove non-tRNA reads and unmature-tRNA reads respectively.

After this first mapping step, a second library is generated by adding 3’ CCA tails and removing introns from tRNA genes. All 100% identical sequences of this so-called *mature* tRNAs are clustered to avoid redundancy. Next, the subset of filtered reads from the first mapping is aligned against the clustered mature tRNAs using Segemehl [v0.3.1]^59^. Mapped reads are then realigned with GATK IndelRealigner [v3.8]^60^ to reduce the number of mismatching bases across all reads.

For quantification, isoacceptors were quantified as reads per million (RPM). In order to increase the coverage for anticodon-level quantification, we consider all reads that map unambiguously to a certain isoacceptor, even though they ambiguously map to different isodecoders (i.e. tRNA genes that differ in their sequence but share the same anticodon). Ambiguous reads mapping to genes of different isoacceptors were discarded.

#### Relative Codon Usage (RCU)

The RCU is defined as the contribution of a certain codon to the amino acid it belongs to. The RCU of all synonymous codons therefore sum up to 1.

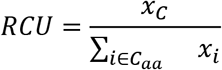

where *x_C_* refers to the abundance of the codon *C*, and *C_aa_* is the set of all synonymous codons.

#### Relative tRNA Adaptation Index (RtAI)

As described by dos Reis et al. (2003, 2004), the tAI weights every codon based on the wobble-base codon-anticodon interaction rules. Let *c* be a codon, then the decoding weight is a weighted sum of the square-root-normalized tRNA abundances *tRNA_cj_* for all tRNA isoacceptors *j* that bind with affinity (1 - *s_cj_*) given the wobble-base pairing rules *n_c_*. However, while dos Reis et al., (2004) assumes that highly expressed genes are codon-optimized, here we use the non-optimized s-values to avoid a circularity in our reasoning:

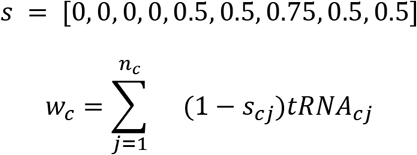

For better comparison with the SDA, an amino-acid-normalized tAI measure is defined by dividing each tAI weight by the maximum weight among all codons within each amino acid family.

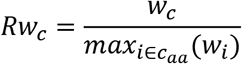

And therefore the RtAI of a certain protein is the product of weights *Rw* of each codon *i_k_* at the triplet position *k* throughout the full gene length *l_g_*, and normalized by the length.

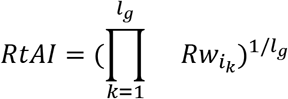

#### Supply-to-Demand Adaptation (SDA)

The SDA aims to consider not only tRNA abundances, but also the codon usage demand. In doing so, it constitutes a global measure of translation control, since the efficiency of a certain codon depends both on its complementary anticodon abundance as well as the demand for such anticodon by other transcripts. This global control has been indeed established to play an important role in defining optimal translation programs^63^.

The definition of the SDA is based on similar previously published metrics^2,64,65^, which consists of a ratio between the anticodon supply and demand. On the one hand, the anticodon supply is defined as the relative tAI weights *Rw* (see previous section). On the other, the anticodon demand is estimated from the codon usage at the transcriptome level. It is computed as the frequency of each codon in a transcript weighted by the corresponding transcript expression, and finally summing up over all transcripts. Let *c* be a codon, then the codon usage is a weighted sum of the counts of codon *c_i_* in gene *j* weighted by the mRNA-seq abundance *mRNA_j_* for all genes in the genome *g:*

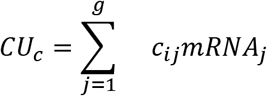

Similarly to the supply, the anticodon demand is then normalized within each amino acid family:

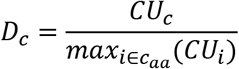

Finally, the SDA weights (SDAw) are defined as the ratio between the codon supply *S_c_* and demand *D_c_*:

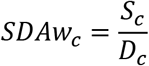

And therefore the SDA of a certain protein is the product of weights *SDAw* of each codon *i_k_* at the triplet position *k* throughout the full gene length *l_g_*, and normalized by the length.

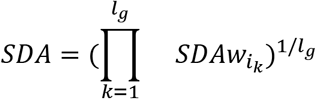

#### Internal clustering validity

Three indexes were used to determine the clustering performance of the RCUs based on different viral features. These are “internal” metrics, since they evaluate the quality of a certain grouping using measures of the dataset itself (homogeneity of clusters, distances within and between clusters, etc.).

- WB index is a ratio of the sum-of-squares (SS) within clusters and the SS between clusters, normalized by the number of clusters^66^. Therefore, low values of the WB index are indicative of good clustering.
- Dunn index considers the inter-cluster distance and diameter of the cluster hypersphere^67^. A higher Dunn index indicates better clustering.
- Silhouette Coefficient ranges from −1 to +1, and measures how similar an object is to its own cluster (intra-cluster distance) compared to other clusters (nearest-cluster distance)^68^. A high value indicates a correct clustering.

#### Linear Discriminant Analysis of viral RCU

We applied a Linear Discriminant Analysis (LDA) to the viral RCUs, taking for each virus the average RCU of its proteins. We assigned each virus to its corresponding tropism (Sup. Table 1) in order to find the linear combination of codon features that maximized differences between viral target tissues. Given the collinear nature of RCUs by definition, the estimated coefficients are impossible to interpret, although it does not hamper the classification performance.

#### Random Forest Classifier

To evaluate the adaptation of the viral proteins to the SDAw of human tissues, we computed their average SDA to each of the 23 TCGA tissues (Sup. Table 3). Using the set of 182 tropism-defined viruses, we had a total of 2891 viral proteins. Taking the 23 tissue-specific SDAs as features, we applied a Random Forest (RF) Classifier, populated with 100 decision trees, using the *scikit-learn* package^69^. Therefore, for each of the six viral tropisms, we developed a model for predicting the tropism-positive versus tropism-negative proteins based on the translational adaptation across tissues. Given that the size of the tropism-positive and tropism-negative groups were often unbalanced, we iteratively sampled equal-sized groups, for n=100 iterations. Furthermore, we validated the results with a stratified 5-fold crossvalidation.

In order to evaluate the performance of the RF models, we computed the Area Under the Curve (AUC) of Receiver Operating Characteristic (ROC) and Precision-Recall (PR) plots (Fig. 2A). We took the average and standard deviation across all iterations. Similarly, we computed the relative feature weights corresponding to each of the 23 TCGA tissues (Fig. 2B).

#### Linear Discriminant Analysis of tissue-specific SDAs

Similar to the RF classifier, we also computed the average proteome SDA per virus in each of the 23 tissues. We then applied a Linear Discriminant Analysis (LDA) to these averaged SDAs. We assigned each virus to its corresponding tropism (Sup. Table 1) in order to find the linear combination of tissue adaptation features that maximized differences between viral target tissues (Extended Data Fig. 1, Sup. Table 3).

#### Gene Set Enrichment Analysis (GSEA)

We analyzed the enrichment of gene sets of the Virus Orthologous Groups using the GSEA algorithm^70^. The score used to generate the ranked list input is specified in the text.

### STATISTICAL ANALYSIS

All details of the statistical analyses can be found in the figure legends. For hypothesis testing, a Wilcoxon rank-sum test was performed. We used a significance value of 0.05.

### DATA AND CODE AVAILABILITY

The code used in this study is available at GitHub (https://github.com/hexavier/tRNA_viruses, https://github.com/hexavier/tRNA_mapping). The published article includes all datasets generated or analyzed during this study.

## ACKNOWLEDGEMENTS

We thank Hannah Benisty and Samuel Miravet-Verde for stimulating and critical discussions. The results published here are in part based on data generated by the TCGA Research Network: https://www.cancer.gov/tcga. We acknowledge the support of the Spanish Ministry of Science and Innovation (MICINN) (PGC2018-101271-B-I00 Plan Estatal), ‘Centro de Excelencia Severo Ochoa’, the CERCA Programme/Generalitat de Catalunya, and the Spanish Ministry of Science and Innovation (MICINN) to the EMBL partnership. The work of X.H. has been supported by a PhD fellowship from the Fundación Ramón Areces.

## AUTHOR CONTRIBUTIONS

Conceptualization, X.H., M.H.S. and L.S.; Methodology, X.H., M.H.S., L.S.; Software, X.H.; Investigation, X.H.; Validation, X.H., M.H.S.; Formal analysis, X.H.; Writing-Original Draft, X.H.; Writing-Review & Editing, X.H., M.H.S., L.S.; Visualisation: X.H., M.H.S., L.S.; Funding Acquisition, L.S.; Supervision, M.H.S. and L.S.

## ADDITIONAL INFORMATION

Further information and requests for resources and reagents should be directed to and will be fulfilled by the corresponding authors L.S. and M.H.S.

## EXTENDED DATA FIGURE LEGENDS

**Extended Data Figure 1.**
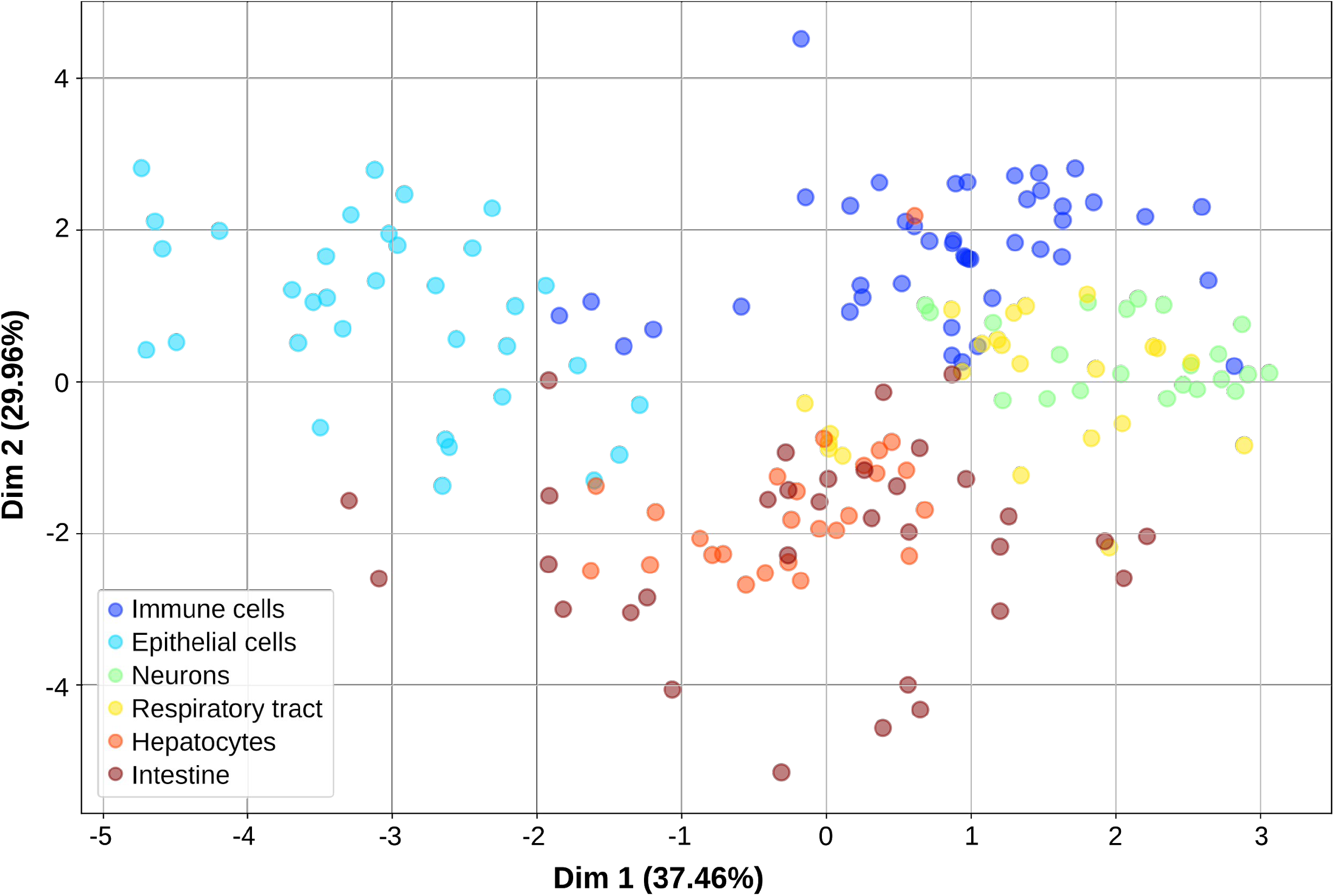
Viruses are adapted to the tRNA-based translational efficiencies of their target tissues, related to Fig. 2. Linear Discriminant Analysis of the Supply-to-Demand Adaptation of the 182 tropism-defined viruses to each of the 23 TCGA tissues, averaged among all the viral proteins and TCGA samples. The features describing the LDA components are available in Sup. Table 3. In brackets, the percentage of variance explained by each of the components.

**Extended Data Figure 2.**
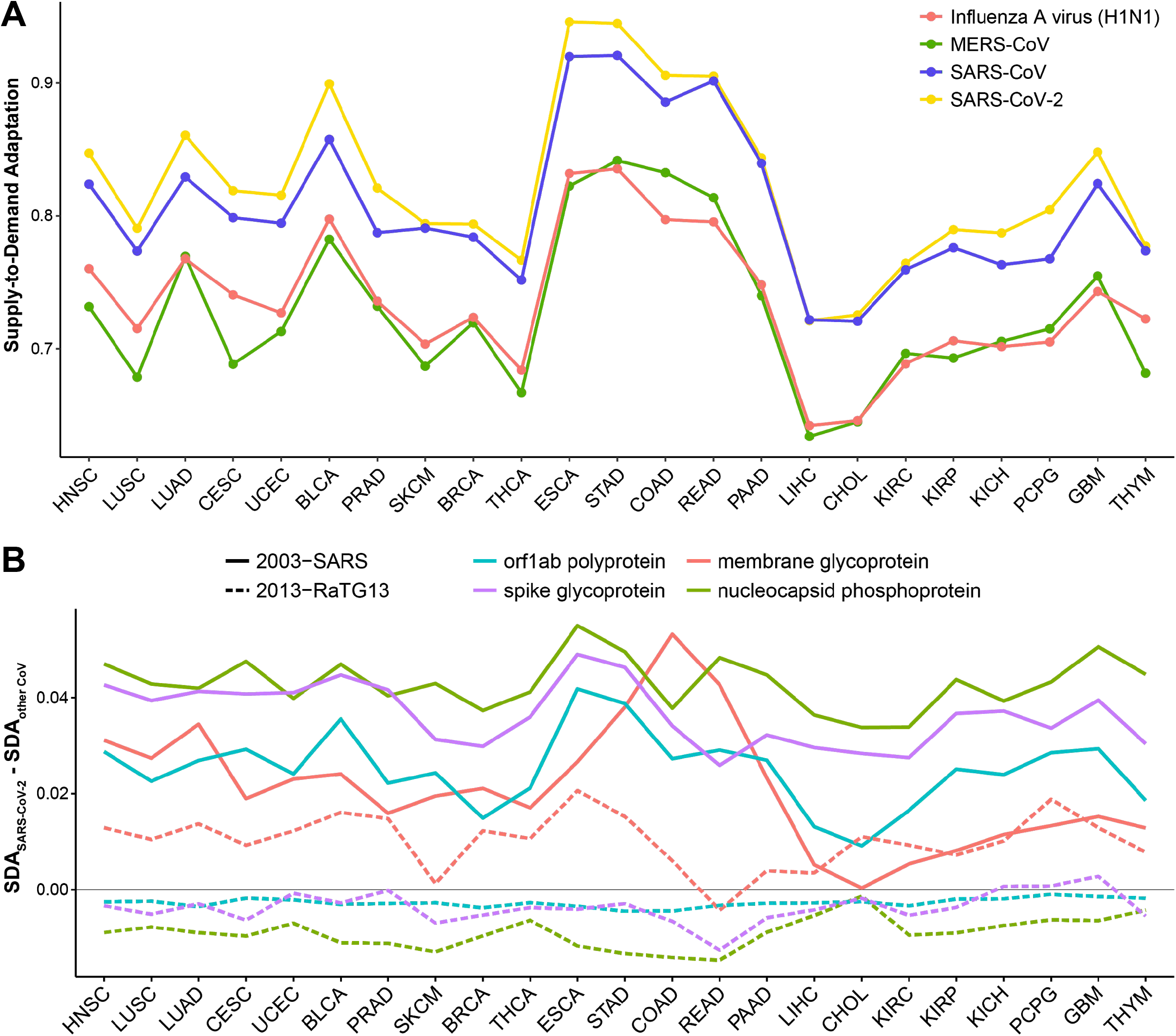
Translational adaptation of SARS-CoV-2 and related viruses among human tissues, related to Fig. 3. (A) Using the codon adaptation weights of all TCGA samples, we compute the average SDA of proteins from coronaviruses and influenza A virus. (B) Difference between the SDA of SARS-CoV-2 and the closest phylogenetically related RaTG13 coronavirus (from bat) or the closest human coronavirus (SARS-CoV). For each strain, the SDA of four of the proteins are shown. Refer to Sup. Table 2 for full TCGA cancer type names.

## SUPPLEMENTARY TABLES

**Sup. Table 1. Human-infecting viruses, related to Fig. 1**. List of all viruses included in this study, together with their related information (genetic material, Baltimore classification, genus, family, tropism). Codon usage and annotation of all proteins of human-infecting viruses.

**Sup. Table 2. TCGA samples, related to Fig. 2–3**. Number and abbreviations of TCGA samples covering 23 cancer types. NT and TP correspond to “Normal Tissue” and “Tumor Primary” respectively.

**Sup. Table 3. Random Forest Dataset, related to Fig. 2**. SDA of all viral proteins across the healthy samples of all TCGA tissues. Feature weights describing the components of the Linear Discriminant Analysis (Extended Data Fig. 1).

**Sup. Table 4. Translational adaptation of SARS-CoV-2, related to Fig. 3**. SDA of viral proteins across the healthy samples of all TCGA tissues, including MERS-CoV, SARS-CoV, SARS-CoV-2 and Influenza A virus. RSEM expression of receptors and related proteins for the entry of MERS-CoV, SARS-CoV, SARS-CoV-2 and Influenza A virus.

**Sup. Table 5. Gene Set Enrichment Analysis of tropism SDA, related to Fig. 4**. Defined correspondences between matching tissues to viral tropisms. Supply-to-Demand Adaptation (SDA) of all tropism-defined viral proteins, with regard to their respective tissue of infection. Among the SDA of all proteins, the Gene Set Enrichment Analysis shows the Virus Orthologous Groups that are better or worse adapted.

**Sup. Table 6. Translational reprogramming upon infection, related to Fig. 4**. tRNA isoacceptor quantification in Reads Per Million (RPM) of all small RNA-seq samples from cell lines upon viral infection. Using their tRNA quantification, the translational efficiency of viral proteins in each sample is estimated using the relative tAI (RtAI).

## REFERENCES

1. Grantham, R., Gautier, C., Gouy, M., Jacobzone, M. & Mercier, R. Codon catalog usage is a genome strategy modulated for gene expressivity. Nucleic Acids Res. 9, r43–74 (1981).

2. Hernandez-Alias, X., Benisty, H., Schaefer, M. H. & Serrano, L. Translational efficiency across healthy and tumor tissues is proliferation-related. Mol. Syst. Biol. 16, e9275 (2020).

3. Plotkin, J. B. & Kudla, G. Synonymous but not the same: the causes and consequences of codon bias. Nat. Rev. Genet. 12, 32–42 (2011).

4. Sharp, P. M., Stenico, M., Peden, J. F. & Lloyd, A. T. Codon usage: mutational bias, translational selection, or both? Biochem. Soc. Trans. 21, 835–841 (1993).

5. Morgado, S. & Vicente, A. C. Global In-Silico Scenario of tRNA Genes and Their Organization in Virus Genomes. Viruses-Basel 11, 180 (2019).

6. Lucks, J. B., Nelson, D. R., Kudla, G. R. & Plotkin, J. B. Genome Landscapes and Bacteriophage Codon Usage. Plos Comput. Biol. 4, e1000001 (2008).

7. Carbone, A. Codon Bias is a Major Factor Explaining Phage Evolution in Translationally Biased Hosts. J. Mol. Evol. 66, 210–223 (2008).

8. Ahlgren, N. A., Ren, J., Lu, Y. Y., Fuhrman, J. A. & Sun, F. Alignment-free d2* oligonucleotide frequency dissimilarity measure improves prediction of hosts from metagenomically-derived viral sequences. Nucleic Acids Res. 45, 39–53 (2017).

9. Ren, J., Ahlgren, N. A., Lu, Y. Y., Fuhrman, J. A. & Sun, F. VirFinder: a novel k-mer based tool for identifying viral sequences from assembled metagenomic data. Microbiome 5, 69 (2017).

10. Bahir, I., Fromer, M., Prat, Y. & Linial, M. Viral adaptation to host: a proteome-based analysis of codon usage and amino acid preferences. Mol. Syst. Biol. 5, (2009).

11. Jenkins, G. M. & Holmes, E. C. The extent of codon usage bias in human RNA viruses and its evolutionary origin. Virus Res. 92, 1–7 (2003).

12. Lauring, A. S., Jones, J. O. & Andino, R. Rationalizing the development of live attenuated virus vaccines. Nat. Biotechnol. 28, 573–579 (2010).

13. Pouyet, F., Mouchiroud, D., Duret, L. & Sémon, M. Recombination, meiotic expression and human codon usage. eLife 6, e27344 (2017).

14. Gingold, H. et al. A Dual Program for Translation Regulation in Cellular Proliferation and Differentiation. Cell 158, 1281–1292 (2014).

15. Zhao, K.-N., Gu, W., Fang, N. X., Saunders, N. A. & Frazer, I. H. Gene codon composition determines differentiation-dependent expression of a viral capsid gene in keratinocytes in vitro and in vivo. Mol. Cell. Biol. 25, 8643–8655 (2005).

16. Zhou, J., Liu, W. J., Peng, S. W., Sun, X. Y. & Frazer, I. Papillomavirus capsid protein expression level depends on the match between codon usage and tRNA availability. J. Virol. 73, 4972–4982 (1999).

17. van Weringh, A. et al. HIV-1 modulates the tRNA pool to improve translation efficiency. Mol. Biol. Evol. 28, 1827–1834 (2011).

18. Li, M. et al. Codon-usage-based inhibition of HIV protein synthesis by human schlafen 11. Nature 491, 125–128 (2012).

19. B. Miller, J., Hippen, A. A., M. Wright, S., Morris, C. & G. Ridge, P. Human viruses have codon usage biases that match highly expressed proteins in the tissues they infect. Biomed. Genet. Genomics 2, (2017).

20. World Health Organization. Novel Coronavirus (2019-nCoV) situation reports. https://www.who.int/emergencies/diseases/novel-coronavirus-2019/situation-reports.

21. Sungnak, W., Huang, N., Bécavin, C., Berg, M. & Network, H. L. B. SARS-CoV-2 Entry Genes Are Most Highly Expressed in Nasal Goblet and Ciliated Cells within Human Airways. ArXiv200306122 Q-Bio (2020).

22. Zhao, Y. et al. Single-cell RNA expression profiling of ACE2, the putative receptor of Wuhan 2019-nCov. bioRxiv 2020.01.26.919985 (2020) doi:10.1101/2020.01.26.919985.

23. Baig, A. M., Khaleeq, A., Ali, U. & Syeda, H. Evidence of the COVID-19 Virus Targeting the CNS: Tissue Distribution, Host–Virus Interaction, and Proposed Neurotropic Mechanisms. ACS Chem. Neurosci. (2020) doi:10.1021/acschemneuro.0c00122.

24. Zhang, H. et al. The digestive system is a potential route of 2019-nCov infection: a bioinformatics analysis based on single-cell transcriptomes. bioRxiv 2020.01.30.927806 (2020) doi:10.1101/2020.01.30.927806.

25. Hulo, C. et al. ViralZone: a knowledge resource to understand virus diversity. Nucleic Acids Res. 39, D576–582 (2011).

26. Aiewsakun, P. & Simmonds, P. The genomic underpinnings of eukaryotic virus taxonomy: creating a sequence-based framework for family-level virus classification. Microbiome 6, 38 (2018).

27. Gogakos, T. et al. Characterizing Expression and Processing of Precursor and Mature Human tRNAs by Hydro-tRNAseq and PAR-CLIP. Cell Rep. 20, 1463–1475 (2017).

28. Zhang, Z. et al. Global analysis of tRNA and translation factor expression reveals a dynamic landscape of translational regulation in human cancers. Commun. Biol. 1, 234 (2018).

29. Belalov, I. S. & Lukashev, A. N. Causes and Implications of Codon Usage Bias in RNA Viruses. PLoS ONE 8, e56642 (2013).

30. Shackelton, L. A., Parrish, C. R. & Holmes, E. C. Evolutionary basis of codon usage and nucleotide composition bias in vertebrate DNA viruses. J. Mol. Evol. 62, 551–563 (2006).

31. Drosten, C. et al. Identification of a novel coronavirus in patients with severe acute respiratory syndrome. N. Engl. J. Med. 348, 1967–1976 (2003).

32. Zaki, A. M., van Boheemen, S., Bestebroer, T. M., Osterhaus, A. D. M. E. & Fouchier, R. A. M. Isolation of a novel coronavirus from a man with pneumonia in Saudi Arabia. N. Engl. J. Med. 367, 1814–1820 (2012).

33. Zhou, P. et al. A pneumonia outbreak associated with a new coronavirus of probable bat origin. Nature 579, 270–273 (2020).

34. Broszeit, F. et al. N-Glycolylneuraminic Acid as a Receptor for Influenza A Viruses. Cell Rep. 27, 3284–3294.e6 (2019).

35. Raj, V. S. et al. Dipeptidyl peptidase 4 is a functional receptor for the emerging human coronavirus-EMC. Nature 495, 251–254 (2013).

36. Hoffmann, M. et al. SARS-CoV-2 Cell Entry Depends on ACE2 and TMPRSS2 and Is Blocked by a Clinically Proven Protease Inhibitor. Cell 0, (2020).

37. Wang, K. et al. SARS-CoV-2 invades host cells via a novel route: CD147-spike protein. bioRxiv 2020.03.14.988345 (2020) doi:10.1101/2020.03.14.988345.

38. Lv, L., Li, G., Chen, J., Liang, X. & Li, Y. Comparative genomic analysis revealed specific mutation pattern between human coronavirus SARS-CoV-2 and Bat-SARSr-CoV RaTG13. bioRxiv 2020.02.27.969006 (2020) doi:10.1101/2020.02.27.969006.

39. Knipe, D. M. & Howley, P. M. Fields Virology. (Lippincott Williams&Wilki, 2013).

40. Chang, S. T. et al. Next-generation sequencing of small RNAs from HIV-infected cells identifies phased microrna expression patterns and candidate novel microRNAs differentially expressed upon infection. mBio 4, e00549–00512 (2013).

41. Shi, J. et al. Deep RNA Sequencing Reveals a Repertoire of Human Fibroblast Circular RNAs Associated with Cellular Responses to Herpes Simplex Virus 1 Infection. Cell. Physiol. Biochem. Int. J. Exp. Cell. Physiol. Biochem. Pharmacol. 47, 2031–2045 (2018).

42. Stark, T. J., Arnold, J. D., Spector, D. H. & Yeo, G. W. High-resolution profiling and analysis of viral and host small RNAs during human cytomegalovirus infection. J. Virol. 86, 226–235 (2012).

43. Franzo, G., Tucciarone, C. M., Cecchinato, M. & Drigo, M. Canine parvovirus type 2 (CPV-2) and Feline panleukopenia virus (FPV) codon bias analysis reveals a progressive adaptation to the new niche after the host jump. Mol. Phylogenet. Evol. 114, 82–92 (2017).

44. Luo, W. et al. The fit of codon usage of human-isolated avian influenza A viruses to human. Infect. Genet. Evol. 104181 (2020) doi:10.1016/j.meegid.2020.104181.

45. Wong, E. H. M., Smith, D. K., Rabadan, R., Peiris, M. & Poon, L. L. M. Codon usage bias and the evolution of influenza A viruses. Codon Usage Biases of Influenza Virus. BMC Evol. Biol. 10, 253 (2010).

46. Pan, L. et al. Clinical characteristics of COVID-19 patients with digestive symptoms in Hubei, China: a descriptive, cross-sectional, multicenter study. 25.

47. Zhou, Z. et al. Effect of gastrointestinal symptoms on patients infected with COVID-19. Gastroenterology 0, (2020).

48. Li, Y.-C., Bai, W.-Z. & Hashikawa, T. The neuroinvasive potential of SARS-CoV2 may play a role in the respiratory failure of COVID-19 patients. J. Med. Virol. n/a,.

49. Mao, L. et al. Neurological Manifestations of Hospitalized Patients with COVID-19 in Wuhan, China: a retrospective case series study. medRxiv 2020.02.22.20026500 (2020) doi:10.1101/2020.02.22.20026500.

50. Andersen, K. G., Rambaut, A., Lipkin, W. I., Holmes, E. C. & Garry, R. F. The proximal origin of SARS-CoV-2. Nat. Med. 1–3 (2020) doi:10.1038/s41591-020-0820-9.

51. Chen, F. et al. Dissimilation of synonymous codon usage bias in virus-host coevolution due to translational selection. Nat. Ecol. Evol. (2020) doi:10.1038/s41559-020-1124-7.

52. Pavon-Eternod, M. et al. Vaccinia and influenza A viruses select rather than adjust tRNAs to optimize translation. Nucleic Acids Res. 41, 1914–1921 (2013).

53. Mioduser, O., Goz, E. & Tuller, T. Significant differences in terms of codon usage bias between bacteriophage early and late genes: a comparative genomics analysis. Bmc Genomics 18, 866 (2017).

54. Walker, P. J. et al. Changes to virus taxonomy and the International Code of Virus Classification and Nomenclature ratified by the International Committee on Taxonomy of Viruses (2019). Arch. Virol. 164, 2417–2429 (2019).

55. Alexaki, A. et al. Codon and Codon-Pair Usage Tables (CoCoPUTs): Facilitating Genetic Variation Analyses and Recombinant Gene Design. J. Mol. Biol. 431, 2434–2441 (2019).

56. Athey, J. et al. A new and updated resource for codon usage tables. BMC Bioinformatics 18, 391 (2017).

57. Chan, P. P. & Lowe, T. M. tRNAscan-SE: Searching for tRNA Genes in Genomic Sequences. Methods Mol. Biol. Clifton NJ 1962, 1–14 (2019).

58. Hoffmann, A. et al. Accurate mapping of tRNA reads. Bioinformatics 34, 1116–1124 (2018).

59. Hoffmann, S. et al. Fast mapping of short sequences with mismatches, insertions and deletions using index structures. PLoS Comput. Biol. 5, e1000502 (2009).

60. McKenna, A. et al. The Genome Analysis Toolkit: a MapReduce framework for analyzing next-generation DNA sequencing data. Genome Res. 20, 1297–1303 (2010).

61. dos Reis, M., Wernisch, L. & Savva, R. Unexpected correlations between gene expression and codon usage bias from microarray data for the whole Escherichia coli K-12 genome. Nucleic Acids Res. 31, 6976–6985 (2003).

62. dos Reis, M., Savva, R. & Wernisch, L. Solving the riddle of codon usage preferences: a test for translational selection. Nucleic Acids Res. 32, 5036–5044 (2004).

63. Frumkin, I. et al. Codon usage of highly expressed genes affects proteome-wide translation efficiency. Proc. Natl. Acad. Sci. U. S. A. 115, E4940–E4949 (2018).

64. Gingold, H., Dahan, O. & Pilpel, Y. Dynamic changes in translational efficiency are deduced from codon usage of the transcriptome. Nucleic Acids Res. 40, 10053–10063 (2012).

65. Pechmann, S. & Frydman, J. Evolutionary conservation of codon optimality reveals hidden signatures of cotranslational folding. Nat. Struct. Mol. Biol. 20, 237–243 (2013).

66. Zhao, Q. & Fränti, P. WB-index: A sum-of-squares based index for cluster validity. Data Knowl. Eng. 92, 77–89 (2014).

67. Dunn, J. C. Well-Separated Clusters and Optimal Fuzzy Partitions. J. Cybern. 4, 95–104 (1974).

68. Al-Zoubi, M. B. & Raw, M. An Efficient Approach for Computing Silhouette Coefficients. J. Comput. Sci. 4, (2008).

69. Pedregosa, F. et al. Scikit-learn: Machine Learning in Python. J. Mach. Learn. Res. 12, 2825–2830 (2011).

70. Subramanian, A. et al. Gene set enrichment analysis: A knowledge-based approach for interpreting genome-wide expression profiles. Proc. Natl. Acad. Sci. U. S. A. 102, 15545–15550 (2005).

